# Imaging decision-related neural cascades in the human brain

**DOI:** 10.1101/050856

**Authors:** Jordan Muraskin, Truman R. Brown, Jennifer M. Walz, Bryan Conroy, Robin I. Goldman, Paul Sajda

## Abstract

Perceptual decisions depend on coordinated patterns of neural activity cascading across the brain, running in time from stimulus to response and in space from primary sensory regions to the frontal lobe. Measuring this cascade and how it flows through the brain is key to developing an understanding of how our brains function. However observing, let alone understanding, this cascade, particularly in humans, is challenging. Here, we report a significant methodological advance allowing this observation in humans at unprecedented spatiotemporal resolution. We use a novel encoding model to link simultaneously measured electroencephalography (EEG) and functional magnetic resonance imaging (fMRI) signals to infer the high-resolution spatiotemporal brain dynamics taking place during rapid visual perceptual decision-making. After demonstrating the methodology replicates past results, we show that it uncovers a previously unobserved sequential reactivation of a substantial fraction of the pre-response network whose magnitude correlates with decision confidence. Our results illustrate that a temporally coordinated and spatially distributed neural cascade underlies perceptual decision-making, with our methodology illuminating complex brain dynamics that would otherwise be unobservable using conventional fMRI or EEG separately. We expect this methodology to be useful in observing brain dynamics in a wide range of other mental processes.

## Introduction

The detailed spatiotemporal brain dynamics that underlie human decision-making are difficult to measure. Invasive techniques with sufficient temporal or spatial resolution, such as depth electrodes or cortical arrays used with epilepsy patients, are only feasible in rare cases and, in addition, do not capture activity from the entire brain. In comparison, non-invasive measures such as electroencephalography (EEG) and magnetoencephalography (MEG) suffer from poor spatial resolution, and blood oxygen level dependent functional MRI (BOLD fMRI) from poor temporal resolution and indirect coupling to neural activity (e.g. fMRI)^1^. In spite of this, EEG, MEG, and fMRI have been used individually to study perceptual decision-making in the human brain, although, by themselves they provide a limited view of the underlying brain dynamics ^2^.

Recently, methods enabling simultaneous acquisition of EEG and fMRI (EEG/fMRI) have led to varied analytic approaches aimed at integrating the electrophysiological and hemodynamic information contained in the joint measurements. Such approaches offer the potential to provide a comprehensive picture of global brain dynamics, and will likely offer new insights into how the brain makes rapid decisions ^3, 4^. Some of the techniques that have been proposed for combining multi-modal brain signals have separately analyzed the EEG and fMRI data and subsequently juxtaposed the results^5, 6^, while others attempt for a truly integrated approach in order to fully exploit the joint information contained in the data sets ^7^. In general, simultaneous EEG/fMRI and the associated analysis techniques have been used to identify neuronal sources of EEG trial-to-trial variability, linking them to cognitive processes such as attention ^8^ and inhibition ^9^.

Many previous studies have used known EEG markers (P1, N2, N170, P300, α- rhythm) or data driven approaches such as Independent Component Analysis (ICA) to combine EEG with fMRI data ^4, 8–16^. One promising approach has been to use supervised machine-learning techniques (e.g. classifiers) to find relevant projections of the EEG data, where single-trial variability of the electrophysiological response along these projections can be correlated in the fMRI space. Goldman, et al. ^17^, Walz, et al. ^18^ and Fouragnan, et al. ^19^ have demonstrated this technique on visual and auditory paradigms. This methodology has been shown to localize cortical regions that modulate with the task while preserving the temporal progression of task-relevant neural activity.

Here we combine a classification methodology with an encoding model that relates the trial-to-trial variability in the EEG to what is observed in the simultaneously acquired fMRI. Encoding models have become an important machine learning tool for analysis of neuroimaging data, specifically fMRI ^20^. In most cases encoding models have been used to learn brain activity that encodes or represents features of a stimulus, such as visual orientation energy in an image/video ^21–23^, acoustic spectral power in sound/speech ^24^, or visual imagery during sleep ^25^. In the method presented here, we employ an encoding model to directly relate the simultaneously collected data from the two neuroimaging modalities—instead of features derived from the stimulus, they are derived from EEG component trial-to-trial variability. Specifically, we learn an encoding in the spatially precise fMRI data from the temporally precise trial-to-trial variability of EEG activity predictive of the level of stimulus evidence. This approach leverages the fact that the level of stimulus evidence, as measured via EEG, persists across the trial ^26, 27^, and that by discriminating this information in a time-localized way, one can temporally “tag” specific cortical areas by their trial-to-trial variability.

Using our framework for learning the BOLD signal encoding of task-relevant and temporally precise EEG component variability, we unravel the cascade of activity from the representation of sensory input to decision formation, decision action, and decision monitoring. A particularly novel finding is that after the activation of decision monitoring regions (i.e. ACC), we see a reactivation of pre-response networks, where the strength of this reactivation correlates with measures of decision confidence. This specific reactivation, as well as the entire spatio-temporal cascade, is completely unobservable using conventional fMRI-only or EEG-only methodologies.

## Results

In this study, we used a visual alternative forced choice (AFC) task where subjects were shown brief presentations of pictures corrupted by noise and instructed to rapidly discriminate between object categories. On any given trial, the level of noise, or stimulus evidence, was varied randomly. The task itself, as well as similar visual decision-making tasks ^28^, is believed to engage an extensive set of cortical areas in a coordinated fashion, including regions that are responsible for sensory encoding, evidence accumulation, decision formation, and response and decision monitoring. However, the dynamic interplay of these regions has never been observed in humans. Here we exploit previously reported findings regarding the sensitivity of the EEG and fMRI signals to the level of stimulus evidence during a perceptual decision-making task. Specifically, previous work has shown differential neural responses to high vs. low stimulus evidence in trial averaged EEG event-related potentials (ERPs), where this difference persists across the trial^26, 27^. Similarly, fMRI studies have shown that for perceptual decision making tasks a number of spatially-distributed cortical areas significantly correlate with the level of stimulus evidence^29, 30^. We leverage the fact that the level of stimulus evidence is expressed temporally in the EEG and spatially in the fMRI to “tag” voxels with a time. Specifically, using a classification methodology (i.e. discriminative components) we identify temporally precise expressions of the level of stimulus evidence that then can be spatially localized through an encoding model of the fMRI BOLD data.

We collected simultaneous EEG/fMRI data from 21 subjects as they performed a 3-AFC task discriminating between faces, cars, and houses (Fig. 1A). Subjects were instructed to discriminate the object class after briefly viewing an image corrupted by varying levels of noise (Fig. 1B) and respond by pressing one of three buttons. Overall, subjects responded with accuracies of 94 ± 5% and 58 ± 12% and with response times of 634 ± 82ms and 770 ± 99ms for high and low stimulus evidence trials, respectively (Fig. 1 C, D). Subject accuracies and response times across stimulus types (faces, cars, houses) for low stimulus evidence trials were similar; however, for high stimulus-evidence trials subject accuracies were higher and response times were shorter for faces than for cars or houses (See Supplemental Information Fig. S1).

**Figure 1.**
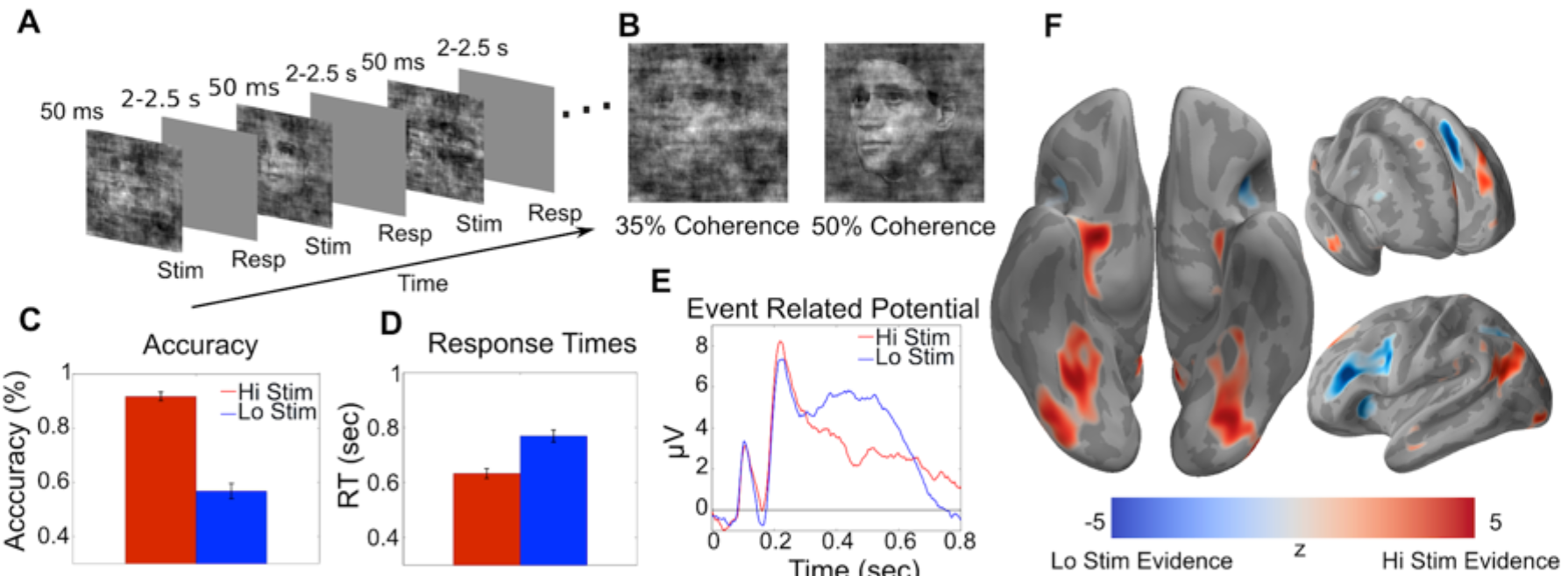
**Paradigm and traditional EEG and fMRI results A**, 3-AFC task where stimulus evidence for each category is modulated by varying the phase coherence in the images. **B**, Example of face images with high stimulus evidence (high coherence: 50%) and low stimulus evidence (low coherence: 35%). **C**, Behavioral performance shows significant differences, as a function of stimulus evidence, in accuracy (p< 10^−12^, paired t-test) and **D**, response time (p< 10^−8^, paired t-test) across the group. **E**, Grand average stimulus-locked event related potentials (ERPs) for electrode Pz show that differences in stimulus evidence span the time from stimulus to response. **F**, fMRI analysis showing cortical areas correlated with high (red) vs. low (blue) stimulus evidence across the entire trial (Z> 2.57 with p< 0.01 Family-Wise Error cluster corrected).

### GLM based analysis of BOLD fMRI shows superposition of cortical areas correlated with stimulus evidence

A traditional general linear model (GLM) analysis of the fMRI (see Methods) revealed differences in BOLD activation between the two stimulus evidence conditions (Fig. 1F, SI Table 1). Brain regions showing greater BOLD activation to high vs. low stimulus evidence trials included areas associated with early visual perception and the default mode network^26^, such as fusiform gyrus, parahippocampal gyrus, lateral occipital cortex, superior frontal gyrus, and posterior cingulate cortex. Regions with greater BOLD activation to low vs. high stimulus evidence trials included areas in the executive control and difficulty networks such as dorsal lateral prefrontal cortex, anterior cingulate cortex, intraparietal sulcus, and insula. Overall, these GLM results for the BOLD data reproduced previous results in the literature where similar stimuli and paradigms were used ^29^(Fig. S2A).

### Extracting temporally localized EEG signatures of stimulus evidence variability

The traditional fMRI results showed multiple brain regions correlated with the difficulty, or stimulus evidence, of the trial; however, this traditional approach does not enable one to infer the relative timing of these fMRI activations. To infer timing at a scale of tens of milliseconds, we used linear classification^31, 32^ of the EEG to extract trial-to-trial variability related to stimulus evidence at specified post-stimulus time points.

The basic idea is illustrated in Figure 2, where hypothetical neural activity is shown for two different regions that are constituents of the perceptual decision-making network. Averaging over trials would clearly reveal a difference in the mean neural activity between high and low stimulus evidence. However, the two regions contribute differentially to the network, with one region encoding the stimulus evidence (Region 1) and the other integrating it over time (Region 2); both are sensitive to the level of stimulus evidence, though varyingly so at different times in the trial. By taking advantage of this sensitivity to the stimulus evidence, we can learn EEG discriminant components, i.e. spatial filters, that best classify trials at different time windows given the neural data. We used the trial-to-trial variability along these component directions as features to uniquely tag fMRI voxels with the specific time window of the component. This tagging is done by building an encoding model of the features, given the BOLD signal, details of which are described in the following section.

**Figure 2.**
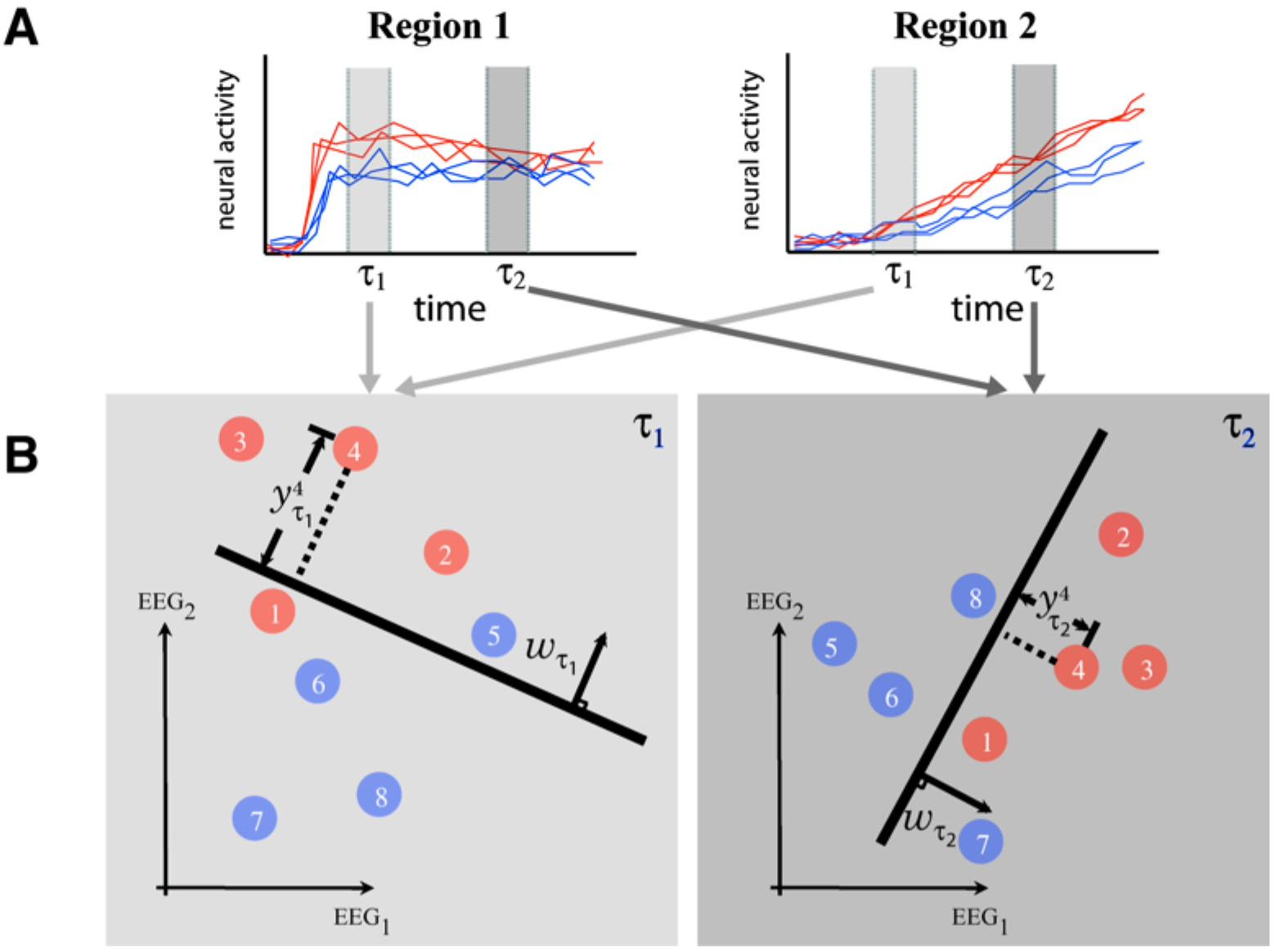
**Temporally precise trial-to-trial EEG variability tags brain regions during decision-making A**, Illustration of how trial-to-trial variability of neural activity in spatially distinct cortical areas can be used to tag brain regions. In this hypothetical example Region 1 is involved in sensory encoding while Region 2 integrates sensory evidence to form a decision (in NHP literature, Region 1 might represent MT, while Region 2 LIP). Neural activity across the trial is shown for two stimulus types, one with high sensory evidence for the choice (red curves) and one with low sensory evidence (blue curves). Also shown are two temporal windows (τ_1_ and τ_2_) that represent different times during the trial. **B**, Linear classifiers are trained to separate trials based on the two levels of stimulus evidence at specific temporal windows. Shown are classifiers (parameterized by weight vectors *w*_1_ and *w*_2_) for two temporal windows (τ_1_ and τ_2_) with respect to two EEG sensors (for simplicity only two dimensions of the full N=43 sensor space are shown. Though the component hyperplane is optimal for the full 43 dimensions, when projected to a line in two dimensions for illustration, it may appear that the separation is sub-optimal). This yields an EEG discriminant component for each temporal window. Variability along these components serves as a unique feature vector for temporally tagging the BOLD data—e.g. variability along an EEG component trained with data from τ_1_ tags BOLD voxels with time τ_1_ while variability along an EEG component trained with data from τ_2_ tags them with τ_2_.

We constructed EEG components by learning linear classifiers at 25ms steps, starting from stimulus onset to 50ms past the average low stimulus evidence response time. We chose a time step of 25ms due to an empirical analysis showing a half width of 50ms in the temporal autocorrelation of the EEG data, though in principle this methodology allows for temporal resolution up to the EEG sampling rate. Each classifier was associated with a set of discriminant values, which can be represented as a vector y_T_; each element of the vector is the distance of a given trial to the discrimination boundary for the classifier at time step τ (Fig. 2). This distance can be interpreted as a measure of the EEG classifier’s estimate of the level of stimulus evidence for that trial^17,18,31–34^.

Results of the EEG analysis show discriminating information for stimulus evidence spanning the trial (see Fig. 4A), beginning roughly 175ms post-stimulus to past the average response times. A dip occurs around 300ms, indicating stimulus evidence is less discriminative at this time and serves to demarcate early and late cognitive processes. The early process corresponded to the time of the D220 ERP component, which has been shown to modulate with the degree of task difficulty, whether via stimulus noise or task demands^35^. The later and more prolonged component is likely related to more complex cognitive and motor preparatory processes that differ between high and low stimulus evidence trials. Importantly, although the early and late EEG components were both discriminative, we found their trial-to-trial variability to be uncorrelated (Figs. 4B and S3E), indicating that while the discriminating information (level of stimulus evidence) persists across the trial, it couples differently to processes across time.

### An encoding model links fMRI activations with temporally distinct EEG trial-to-trial variability

After extracting the trial-to-trial variability from the EEG discriminant components, feature vectors y_τ_ are collected across time steps, τ, along with a response time vector to construct a matrix *Y*. This matrix is the temporally precise representation of the trial-to-trial EEG variability that reflects high vs. low stimulus evidence. An encoding model is then fit, namely a model in which weights are estimated for each time-localized EEG window, to predict the trial-to-trial variability of the BOLD response for each fMRI voxel. Figure 3 shows a schematic of the encoding model framework we used and compares it to a traditional encoding model constructed by using features derived directly from the stimulus. Rather than constructing a map that directly relates each voxel to a type of stimulus feature, such as whether it encodes edges, motion or some semantic concept such as “animal”^21–23, 36–38^, our model is used to construct maps that label voxels by the time window of the variability they encode — i.e. it “tags” each voxel with a “time”, or set of times, when it encodes the variability in the given EEG discriminant component(s).

**Figure 3.**
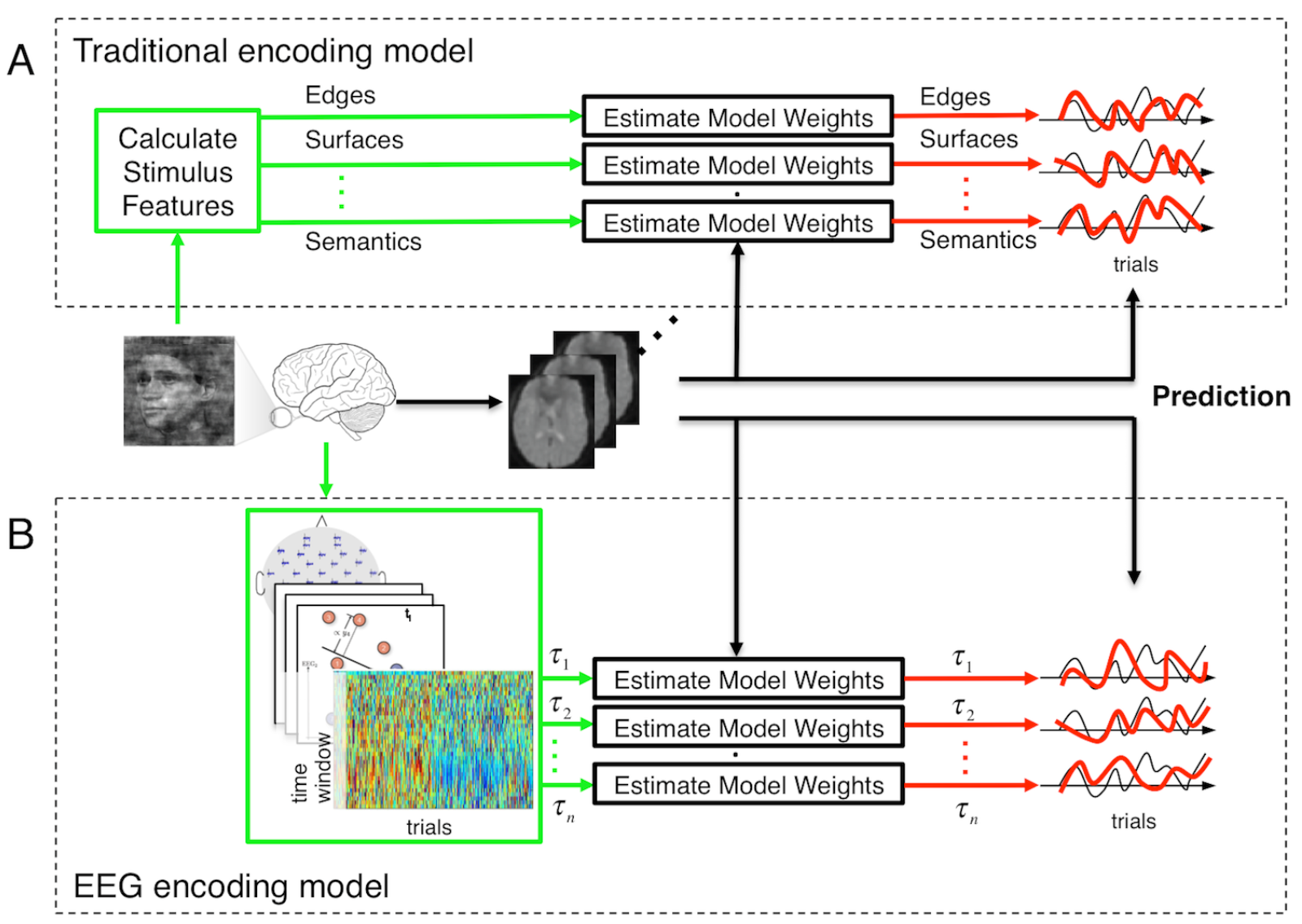
**Encoding models based on stimulus derived features versus EEG variability A**, A traditional encoding model used in fMRI analysis extracts a set of features from the stimulus that are potentially representative of low level structure and high level semantics (green box). Weights are learned to model how these stimulus features are encoded in the fMRI BOLD signal. The resulting encoding model is used to make predictions based on how well different voxels predict the features from novel stimuli. For example, one can create maps of the brain that are labeled based on the stimulus features that each voxel represents. **B**, The same encoding model concept applied to EEG variability (EEG encoding model). Instead of features being estimated from the stimulus, they are derived from EEG component trial-to-trial variability (as in Fig 2a) with each temporal window representing a different feature (green box). Weights are learned so as to model how the EEG variability at a given time window is encoded in the fMRI BOLD. As in the traditional encoding model, predictions on novel stimuli can be done to test the model and results can be used to construct a map—in this case a map of the brain that shows the timing of the EEG component variability that each voxels represents.

It is important to note that this approach does not attempt to improve source localization typically done for EEG/MEG studies. Our approach instead provides the temporal resolution of EEG (ms) and the spatial resolution of fMRI (mm) without the need to solve the ill-posed inverse solution and make the many associated assumptions required for reliable source-localization results^39^.

An example of the quality of the encoding model is shown in Fig. 4C (see also Fig. S2B) where significant voxels from the encoding model are shown in yellow. Fig. 4D shows the trial-to-trial variability of BOLD signal at a specific voxel, comparing it to the variability predicted by the encoding model. Additional validity of the encoding model and single subject results are presented in the Supplemental Information (Fig. S4A/B). The encoding model was also evaluated as a decoding model (see Methods) with the BOLD activity used to predict the trial-to-trial variability in the EEG for unseen data—data on which the encoding model was not trained. Fig. 4E shows these results, expressed as the correlation between the measured and predicted EEG trial-to-trial variability across the 800ms epoch. The shape of the curve is highly consistent with that observed for the EEG data itself (comparing Fig. 4A and Fig. 4E) (additional analysis of the fidelity of the model is provided in the SI, Fig. S3).

**Figure 4.**
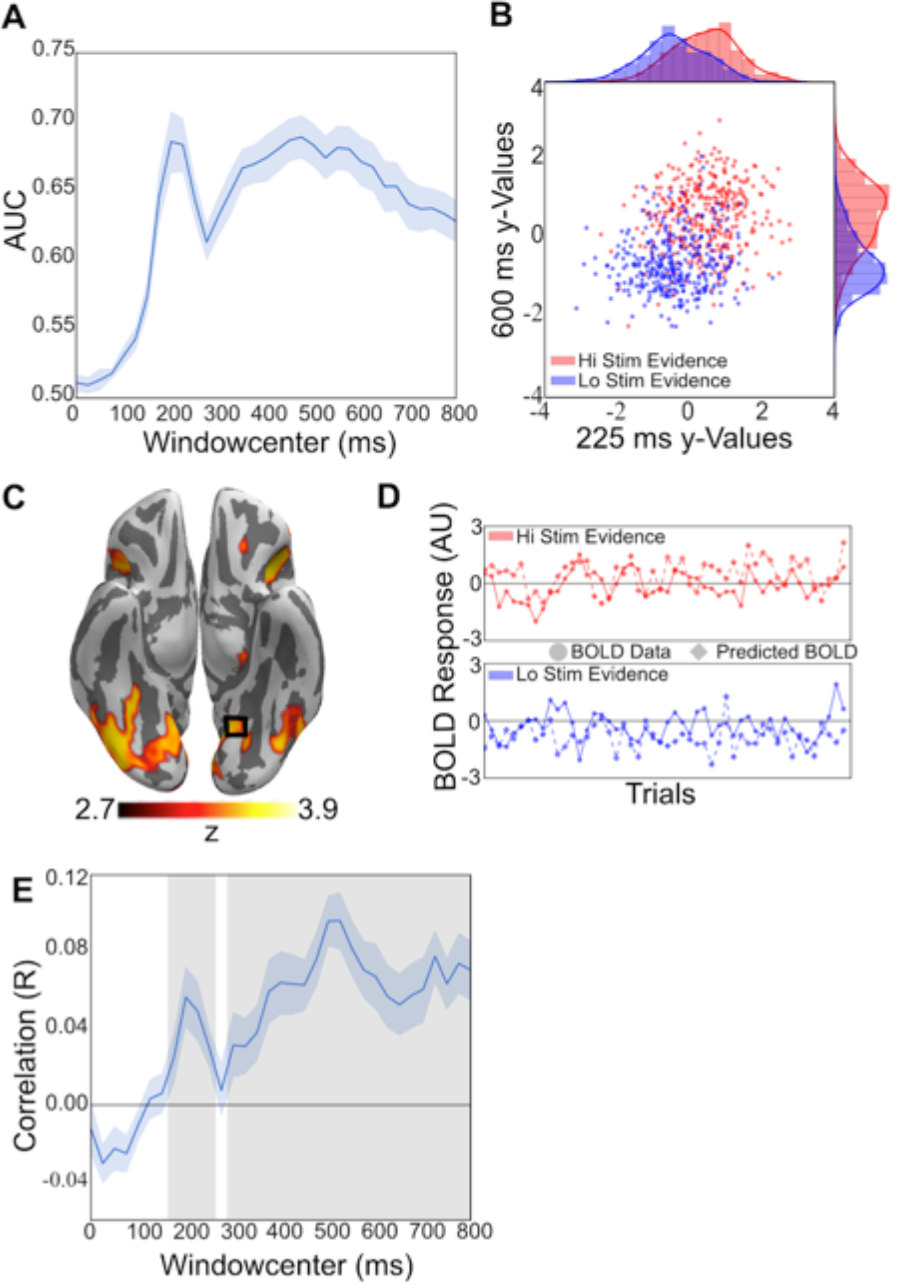
**EEG discrimination and encoding model results A**, Group average area under the receiver operating curve (AUC) for the sliding window logistic regression EEG discrimination analysis, comparing high versus low stimulus evidence trials; standard error across subjects is shown with shading. **B**, A single subject’s discriminating y-value distributions for high (red) and low stimulus evidence (blue) trials for two EEG time points (225ms and 600ms). **C**, Significant fMRI voxels resulting from the group level analysis for the encoding model (p< 0.01 TFCE-False Discovery Rate (FDR) corrected). Activity is seen encompassing early visual processing regions, attention networks, and the task positive network. **D**, A random subset of 100 (50 for each stimulus evidence condition) from 700 total trials of the actual (circle) and predicted (diamond) BOLD responses from the encoding model, for an example subject at a single voxel (MNI X/Y/Zmm: -27/-54/-15, r=0.206, p<10^−6^). High and low stimulus evidence trials are shown separately for clarity. **E**, The averaged correlation of the predicted y-values with the true y-values across the trial duration. Blue shading represents the standard error across subjects. Grey shading indicates significant time windows (p< 0.05 FDR-corrected).

Given the encoding model, we unwrap the BOLD activity across time by identifying weights that are consistent across subjects in space and time (see Methods). Fig. 5 shows these results for a group level analysis. We observe a progression of activity (see Movie S1), at 25ms resolution, which proceeds simultaneously down the dorsal and ventral streams of visual processing for the first 250ms. After that the cascade becomes more complex with activation in the IPS at 425ms and 750ms (see Fig. 6A), reactivation of the SPL at 675ms and activation of ACC at 600ms along with other regions found in the traditional fMRI results. (see Fig. S5, Tables S2 and an additional analysis using dynamic causal modeling ^40^). The reactivation pattern is particularly significant since it would not be observable via a traditional fMRI general linear model (GLM) analysis, which integrates over time and thus superimposes these activities. For example, the changing sign of the middle temporal gyrus (MT) encoding weights in Fig. 6A manifested as no activity in the MT for the traditional fMRI GLM analysis—the change in sign canceled the effective correlation in the GLM (see Fig. 1F and Fig. S1). The areas of activation we find are consistent with previous reports in the literature for human subjects^29, 30^; however, here we are able to link activations across time in a way that was previously only possible with invasive techniques.

**Figure 5.**
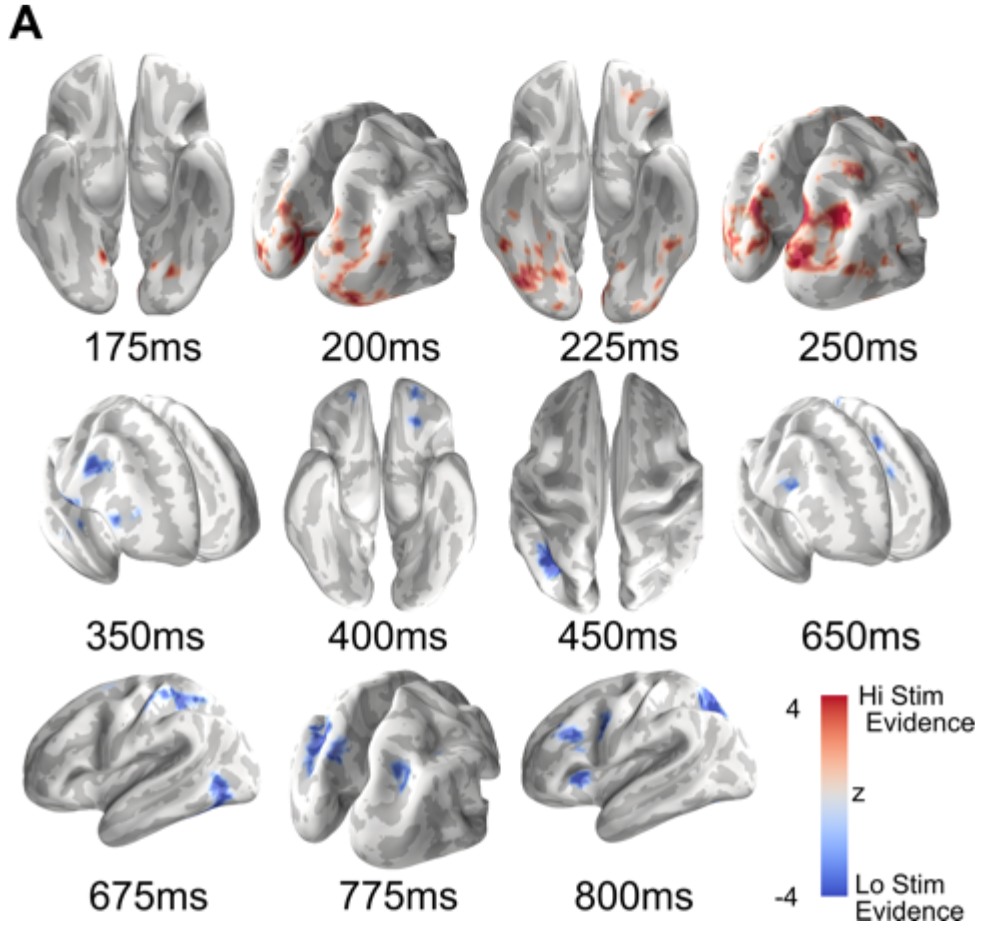
**Group-level encoding model weights results show neural activation cascade** Subset of thresholded (p< 0.05 FDR-Corrected, k=10) group level statistical parametric maps created by stTFCE randomization procedure on the encoding model weight matrices show the progression of spatial activity across the trial. Activation can be seen early in the trial in the occipital regions while progressing more anteriorly later in the trial to executive control areas. Activations in red indicate areas where high stimulus evidence trials had larger activations than low stimulus evidence trials, and blue the inverse.

### Cortical reactivation correlates with decision confidence

Further analysis of the spatiotemporal dynamics (see Fig. 6B), shows that the reactivation pattern in the network occurs after decision-monitoring areas become engaged (i.e. after ACC). Spontaneous reactivation, or “replay”, of neural activity in the human brain has been observed and believed to be important for memory consolidation^41^ and more recently has been hypothesized to play a role in perceptual decision-making by enabling the formation of decision confidence^42^. To test the hypothesis that the reactivation activity we see is in fact related to decision confidence, we used a hierarchical drift diffusion model (DDM)^43, 44^ to fit the behavioral data for high and low stimulus evidence conditions (see Methods). Specifically, our model enables us to define a proxy for decision confidence based on the DDM fits to the behavior^45, 46^. Correlating the reactivation level to this confidence proxy shows a strong and significant monotonic relationship between confidence and the level of reactivation (high stimulus evidence-slope=0.037±0.008, t=4.657, p=3.2×10^−6^; low stimulus evidence-slope=0.062±0.008, t=7.754, p=8.88×10^−15^), with low stimulus evidence trials reactivated more strongly than high stimulus evidence trials (difference in slopes=−0.025±0.011, t=2.189, p=0.029)(see Fig. 7 and Fig. S7). Additionally, reactivation amplitude correlates with behavioral accuracy (Fig. S8) (high stimulus evidence, slope=0.0115±0.0047, t=2.41, p=0.016; low stimulus evidence, slope=0.0104±0.0047, t=2.19, p=0.028). Recursive feature elimination showed that the IPS/SPL and dorsal lateral prefrontal cortex (DLPFC) clusters contributed the most to reactivation/confidence proxy correlation (Fig. 7C).

**Figure 6.**
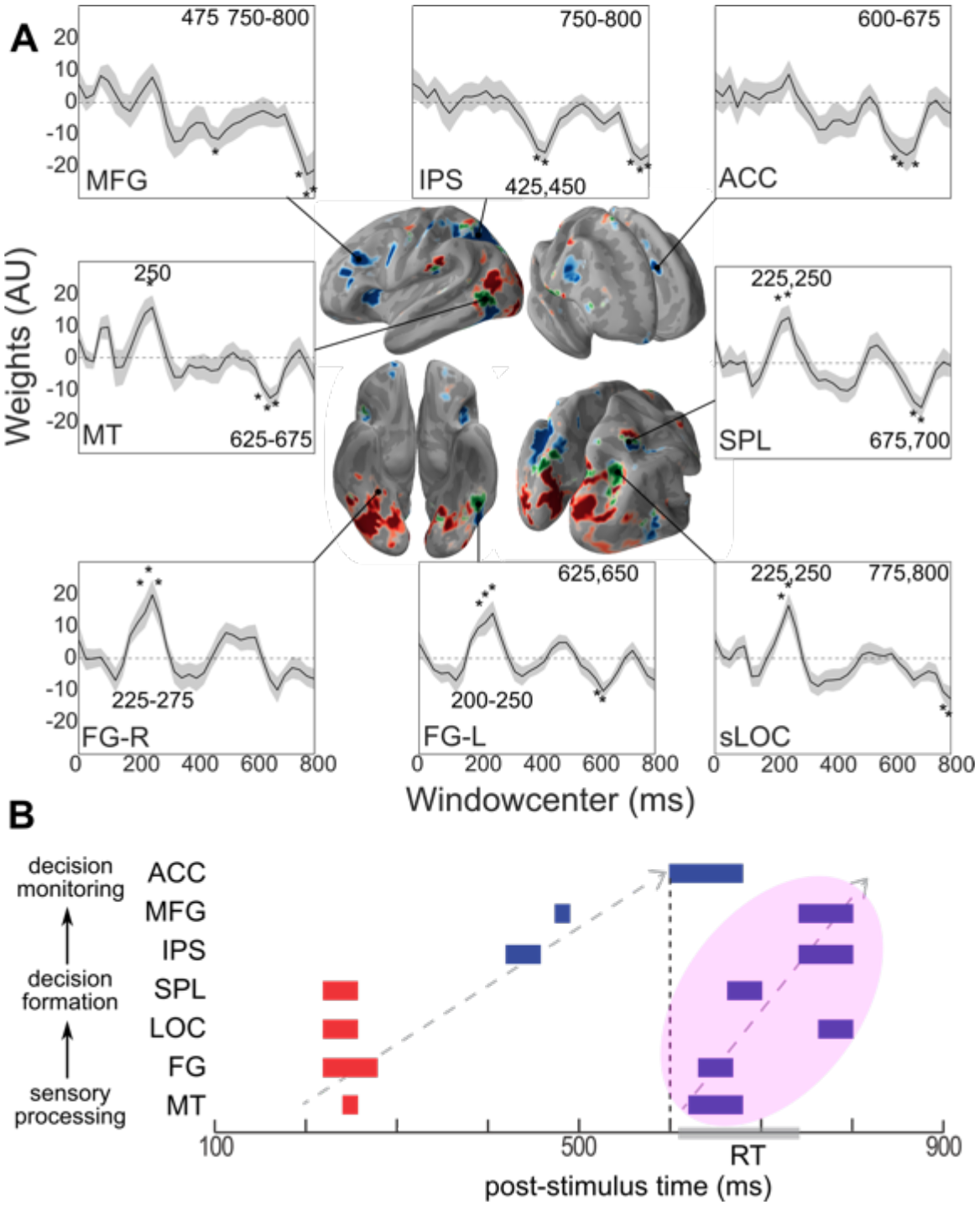
**Spatial-temporal event-related activations show coordinated reactivations. A**, Union across time windows of significant voxels for high (red) and low (blue) stimulus evidence activations. Voxels with activations for both high and low conditions (at different time windows) are displayed in green. Also shown are the encoding model weights for specific voxels, including fusiform gyrus (FG-R):36/-51/-18, (FG-L):-42/- 42/-18, superior lateral occipital cortex (sLOC):24/-63/36, superior parietal lobule (SPL):27/-51/54, anterior cingulate cortex (ACC):-6/24/30, intraparietal sulcus (IPS):- 30/-60/39, middle frontal gyrus (MFG):-45/27/30, middle temporal gyrus (MT):-57/- 60/0. Asterisks indicate significant windows. **B**, Sequence of significant weights showing a “replay” of the network after the onset of ACC activation (shaded ellipse). “Replay” is faster than the initial stimulus driven sequence and strongest for low evidence trials.

**Figure 7.**
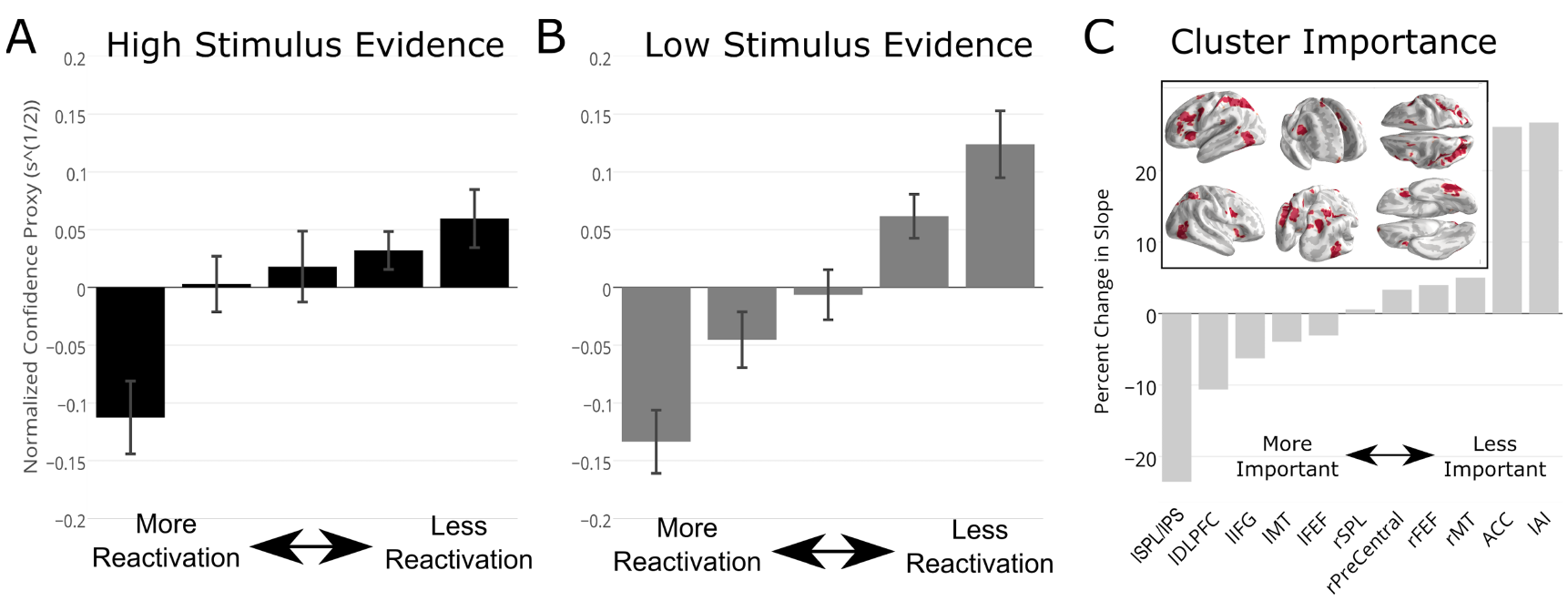
**Trial-to-trial reactivation correlates with decision confidence**. Trial-to-trial reactivation amplitude (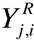 - see Methods) of “replay” correlates with confidence proxy for both high (**A**) and low (**B**) stimulus evidence conditions. Error bars represent standard errors across subjects. (**C**), Stimulus-locked replay activation clusters and feature importance. (**inset**) Regions of interest used in computing the reactivation values for computing confidence proxy correlations. These regions were taken from significant group activations from 600-800ms post stimulus. Regions were then clustered (> 48 voxels) and a secondary analysis for feature importance was performed. Here, we removed each cluster before computing trial-to-trial reactivations and compared the slope of reactivation x confidence proxy when all clusters were present. Panel C shows the ranking of feature importance for each cluster (more negative % change = more importance). Negative changes in slopes show that by removing that cluster the slope of the correlation between reactivation and confidence decreases, indicating the importance of that cluster. Increases in slope indicate that the correlation is higher with that region removed.

## Discussion

We have shown that linking simultaneously acquired EEG and fMRI using a novel encoding model enables imaging of high-resolution spatiotemporal dynamics that underlie rapid perceptual decision-making — decisions made in less than a second. This method, which resolves whole-brain activity with EEG-like temporal resolution, was shown to uncover reactivation processes that would otherwise be masked by the temporal averaging and slow dynamics of traditional fMRI. More broadly, our results demonstrated a general non-invasive data-driven methodology for measuring high spatiotemporal latent neural processes underlying human behavior.

This approach temporally “tags” the BOLD fMRI data by encoding the trial-to-trial variability of the temporally precise task relevant components in simultaneously acquired EEG. In effect, the EEG discrimination indexes the activity of interest at high temporal resolution, defining a feature space, and the trial-to-trial variability of these discriminant components becomes the specific feature values used in the encoding model. For the case presented here, this variability was used to tease apart the cascade of activity modulated by stimulus evidence across the trial, and this allowed us to observe, as never seen before, the spatiotemporal brain dynamics underlying a perceptual decision.

Previous studies have sought to generalize the timing diagram of a perceptual decision through multi-unit recordings in non-human primates^47, 48^ or more broadly in humans^29, 30^ using fMRI. Our results confirmed the general temporal ordering of activations found previously (early visual processing, decision formation, decision monitoring). However, there was a possibility the temporal order we observed using our technique was an artifact of our methodology. To assess this possibility, we performed additional analyses using dynamic causal modeling (DCM) to further validate the temporal activation sequence (see Fig. S6) and show, using a different set of assumptions and method, that the temporal sequence we observe is highly likely under a set of alternative sequences. We found that the most likely model is the one consistent with the time course inferred from our encoding model. The DCM results provide additional evidence that the temporal profile uncovered by the encoding model is a likely temporal decomposition of the superimposed fMRI activations.

The approach we present requires that EEG and BOLD data be collected simultaneously and not in separate sessions in order to exploit the correlations in trial-to-trial variability to “tag” the BOLD data. To show the importance of collecting the data simultaneously, we ran a control analysis that randomly permuted the trials within their stimulus evidence class, thus effectively simulating an EEG and BOLD dataset collected separately. By destroying the link between the EEG and BOLD trials, the encoding model failed to find any consistent activation (Fig. S11/12), indicating the necessity of simultaneous acquisition.

Alternative techniques for fusing simultaneous EEG-fMRI typically do not exploit EEG across the trial and instead only analyze specific ERP components or time windows of interest ^4,8,10,12–19,49,50^. Results from these techniques identify regions that modulate with the specific components, but yield limited information about the timing of other task-relevant regions seen in traditional fMRI contrasts. The methodology developed here extends the work of Goldman, et al. ^17^ and Walz, et al.^18^ by combining their EEG data reduction techniques with techniques developed for encoding stimulus features onto BOLD data^20–23,36,38^, ultimately providing a framework for labeling voxels in task-relevant fMRI contrasts with their timing information (Fig. S2C/E/F).

Clearly, other EEG components that are task-related can be isolated and could potentially be used to “tag” BOLD data. The sliding window linear classification used here acts to reduce the EEG data along a dimension that categorizes stimulus evidence; however, this could be replaced by any other data reduction technique, such as temporally windowed ICA or PCA. Variability along these component directions could then be used in the encoding model to link with the simultaneously collected BOLD data. The choice of data reduction technique (i.e. feature space) would be highly dependent on the nature of the inferences.

Our methodology enabled us to observe reactivation of the pre-response network, spatiotemporal dynamics that would be masked using traditional fMRI analysis.

Interestingly, the reactivation terminated in a network that included the MFG, SPL, and IPS, similar areas previously reported to be reactivated in metacognitive judgments of
confidence in perceptual decisions^42,51,52^. In addition, these areas contributed the most to the correlation to confidence proxy (Fig. 7C). Gherman and Philiastides ^53^ observed this network using a multivariate single-trial EEG approach, coupled with a distributed source reconstruction technique. Fleming, et al. ^42^ and Heereman, et al. ^54^ used BOLD fMRI to show that areas in this network negatively correlate with subjective certainty ratings. Unique to our findings, we saw this reactivation on a single-trial basis after engagement of the ACC, which has been shown to be involved in decision monitoring^53, 55^, and also observed the dynamic sequence leading up to this network reactivation. Our results showed that reactivation/replay occurred on a trial-to-trial basis after a decision, was stronger for difficult decisions, and correlated with decision confidence.

A potential confound in our analysis is that the timing of the reactivation overlaps with some of the response times. To check if the reactivation was pre or post response, we implemented a response-locked encoding model analysis (Fig. S9). The response-locked results showed significant activation pre-response that overlaps with the reactivation network from the stimulus locked analysis. In addition, trial-to-trial reactivation taken from pre-response clusters correlates with confidence proxy similarly to the stimulus locked results (Fig. S10). This provides further evidence that the reactivation is occurring pre-response.

The encoding model we developed was able to decompose traditional fMRI activation maps into their temporal order with significant voxel overlap between the encoding model results and traditional results. The encoding model was also able to show regions that were activated at multiple time points throughout the decision, indicating temporal dynamics that were hidden previously. The regions of activation we found are consistent with earlier findings; however, the work here provided the precise temporal decomposition of these previously reported, temporally superimposed regions of activation. In general, we have shown that simultaneously acquired EEG/fMRI data enables a novel non-invasive approach to visualize high resolution spatial and temporal processing in the human brain with the potential for providing a more comprehensive understanding of the neural basis of complex behaviors.

## Methods

### Subjects

21 subjects (12 male, 9 female; age range 20-35 years) participated in the study. The Columbia University Institutional Review Board (IRB) approved all experiments and informed consent was obtained before the start of each experiment. All subjects had normal or corrected-to-normal vision.

### Stimuli

We used a set of 30 face (from the Max Planck Institute face database), 30 car, and 30 house (obtained from the web) gray scale images (image size 512×512 pixels, 8 bits/pixel). They were all equated for spatial frequency, luminance, and contrast. The stimulus evidence (high or low) of the task was modulated by systematically modifying the salience of the image via randomization of image phase (35% (low) and 50% (high) coherence)^56^.

### Experimental task

The stimuli were used in an event-related three-alternative forced choice (3-AFC) visual discrimination task. On each trial, an image –– either a face, car, or house –– was presented and subjects were instructed to respond with the category of the image by pressing one of three buttons on an MR compatible button controller. Stimuli were presented to subjects using E-Prime software (Psychology Software Tools) and a VisuaStim Digital System (Resonance Technology) with 600x800 goggle display. Over four runs, a total of 720 trials were acquired (240 of each category with 120 high coherence trials) with a random inter-trial interval (ITI) sampled uniformly between 2-2.5s. Each run lasted for 560 seconds.

### JMRI acquisition

Blood-oxygenation-level-dependent (BOLD) T2*-weighted functional images were acquired on a 3T Philips Achieva scanner using a gradient-echo echo-planar imaging (EPI) pulse sequence with the following parameters: Repetition time (TR) 2000ms, echo time (TE) 25ms, flip angle 90°, slice thickness 3mm, interslice gap 1mm, in-plane resolution 3×3 mm, 27 slices per volume, 280 volumes. For all of the participants, we also acquired a standard T1-weighted structural MRI scan (SPGR, resolution 1×1×1mm).

### EEG acquisition

We simultaneously and continuously recorded EEG using a custom-built MR-compatible EEG system^57, 58^, with differential amplifiers and bipolar EEG montage. The caps were configured with 36 Ag/AgCl electrodes including left and right mastoids, arranged as 43 bipolar pairs. Bipolar pair leads were twisted to minimize inductive pickup from the magnetic gradient pulses and subject head motion in the main magnetic field. This oversampling of electrodes ensured data from a complete set of electrodes even in instances when discarding noisy channels was necessary. To enable removal of gradient artifacts in our offline preprocessing, we synchronized the EEG with the scanner clock by sending a transistor-transistor logic pulse at the start of each image volume. All electrode impedances were kept below 20 kΩ, which included 10 kΩ resistors built into each electrode for subject safety.

### Functional image pre-processing

Image preprocessing was performed with FSL (www.fmrib.ox.ac.uk/fsl/). Functional images were spatially realigned to the middle image in the times series (motion-correction), corrected for slice time acquisition, spatially smoothed with a 6mm FWHM Gaussian kernel, and high pass filtered (100s). The structural images were segmented (into grey matter, white matter and cerebro-spinal fluid), bias corrected and spatially normalized to the MNI template using ‘FAST’ ^59^. Functional images were registered into MNI space using boundary based registration (BBR)^60^.

### EEG data preprocessing

We performed standard EEG preprocessing offline using MATLAB (MathWorks) with the following digital Butterworth filters: 0.5 Hz high pass to remove direct current drift,
60 and 120 Hz notches to remove electrical line noise and its first harmonic, and 100 Hz low pass to remove high-frequency artifacts not associated with neurophysiological processes. These filters were applied together in the form of a zero-phase finite impulse response filter to avoid distortions caused by phase delays. We extracted stimulus-locked 1500 ms epochs (-500:1000) and subtracted the mean baseline —200 ms to stimulus onset - from the rest of the epoch. Through visual inspection, we discarded trials containing motion and/or blink artifacts, evidenced by sudden high-amplitude deflections.

### Sliding window logistic regression

We used linear discrimination to associate each trial with the level of stimulus evidence represented in the EEG. We considered high stimulus and low stimulus evidence trials irrespective of behavioral accuracy. Regularized logistic regression was used as a classifier to find an optimal projection for discriminating between high and low stimulus evidence trials over a specific temporal window. A sweep of the regularization parameters was implemented using FaSTGLZ^61^. This approach has been previously applied to identify neural components underlying rapid perceptual decision-making^17,18,31,33,34,45,50,62^.

Specifically, we defined 50ms duration training windows centered at time, τ, ranging from stimulus onset to 800ms following the stimulus in 25ms steps. We used logistic regression to estimate a spatial weighting, on N EEG channels, vector (w_τ_ which is N × 1) that maximally discriminated between EEG sensor array signals E for each class (e.g., high vs. low stimulus evidence trials):

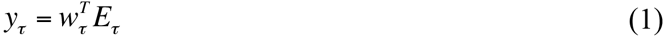

In eqn. 1, E_τ_ is an N × p vector (N sensors per time window τ by p trials). For our experiments, the center of the window (τ) was varied across the trial in 25ms time-steps. We quantified the performance of the linear discriminator by the area under the receiver operator characteristic (ROC) curve, referred to here as AUC, using a leave-one-out procedure. We used the ROC AUC metric to characterize the discrimination performance as a function of sliding our training window (i.e., varying τ). For each subject, this produced a matrix Y where the rows corresponded to trials and the columns to training windows, i.e. Y is the combination of the calculated y_τ_ for each time window.

### Traditional fMRI analysis

We first ran a traditional general linear model (GLM) fMRI analysis in FSL, using event-related (high and low stimulus evidence) and response time (RT) variability regressors. The event-related regressors comprised boxcar functions with unit amplitude and onset and offset matching that of the stimuli. RT variability was modeled using the z-scored RT as the amplitude of the boxcars with onset and offset matching that of the stimulus, and these were orthogonalized to the event-related regressors.

Orthogonalization was implemented using the Gram-Schmidt procedure^63^ to decorrelate the RT regressor from all other event-related regressors. All regressors were convolved with the canonical hemodynamic response function (HRF), and temporal derivatives were included as confounds of no interest. An event-related high versus low stimulus evidence contrast was also constructed. A fixed-effects model was used to model activations across runs, and a mixed-effects approach was used to compute the contrasts across subjects. Activated regions that passed a family-wise error (FWE) ^64^ corrected cluster threshold of p < 0.01 at a z-score threshold of 2.57 were considered significant.

### fMRI deconvolution

Associating fMRI data to each trial is challenging for two main reasons: (a) the temporal dynamics of the hemodynamic response function (HRF) evolve over a longer time-scale than the mean ITI of the event-related design, resulting in overlapping responses between adjacent trials; and (b) the ITI was random for each trial so that the fMRI data was not acquired at a common lag relative to stimulus onset. To overcome these issues, we employed the ‘least squares - separate’ (LS-S) deconvolution^65^ method to estimate the voxel activations for each trial. For every trial, the time series of each voxel was regressed against a “signal” regressor and a “noise” regressor. The “signal” regressor was the modeled HRF response due to that trial (a delta function centered at stimulus onset convolved with a canonical HRF), while the “noise” regressor was the modeled HRF response due to all other trials (superimposed linearly). The resulting regression coefficients of the “signal” regressor represented the estimated voxel activations due to that trial. These voxel activations were then organized into a single brain volume per trial. We extracted 58697 voxels from a common gray matter group mask at 3 mm^3^ spatial resolution that excluded white matter and CSF and assembled the resulting voxel activations into rows of the data matrix F.

### Single subject encoding model

All encoding model analyses were performed in MATLAB. To relate the EEG data with the fMRI, we devised a subject-wise spatio-temporal decomposition using singular value decomposition (SVD). Let F be an m × p matrix denoting m-voxels and p-trials that is the deconvolved high and low stimulus evidence fMRI data for each trial. Let Y be the r x p matrix denoting r-windows (33 EEG_τ_ windows and response time (RT)) and p-trials. For each trial, the first row of Y is the response times while subsequent rows are the y values at each window time. Let W be an m x r matrix that is the weights on Y that solve for F.

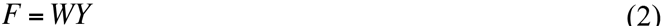

Normally, if we solve for W using the least squares approach, we get:

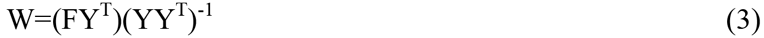

However, each time point might be highly correlated with its neighbors, which reduces the stability of the least-squares regression. We can use SVD to reduce the feature space and improve our estimation of W (the weights on each window). Then for a leave-one-out cross validation, we hold out a single trial from the EEG Y matrix and the corresponding volume from the fMRI data F and train on the remaining trials. We repeated this for all trials.

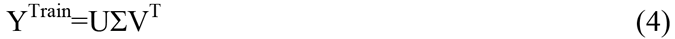

Where U is an r × r orthonormal matrix, ∑ is a r × p diagonal matrix and V is a p × p orthonormal matrix. After SVD on Y^Train^, we reduced the feature dimensions on Y^Train^ to retain 75% of the variance by only keeping v components. To do this, we selected the first v rows of ∑ and zeroed the other rows. We now have 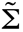 as our reduced spaced matrix. If we now recalculate our least squares solution where we have replaced Y by its reduced form 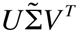 in equation 3:

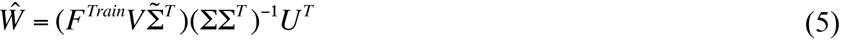

So for each leave one out fold, we first calculated the SVD of the training set. We then calculated the number of components to keep and then solve for *Ŵ*, the weight estimate per fold. To test, we then applied the weights to the left-out test data Y^Test^ to estimate the encoded fMRI data 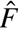 for the encoding part:

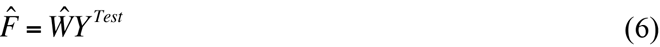

While for the decoding model using the left out test data F^Test^:

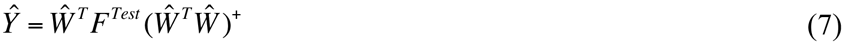

Here, *Ŵ^T^Ŵ* is not invertible, and so we used the pseudo-inverse.

At this point, we have 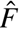, a m x p matrix with m voxels by p trials. For each voxel j, we calculated the correlation of 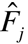 with F_j_, resulting in the matrices R^fMRI^ (Pearson Correlation Map) and P^fMRI^ (p-value map of the Pearson Correlation) that are m x 1. The P^fMRI^ was then converted to a z-score map. We constructed the m x r weight matrix W by taking the average of all the trained Ŵ matrices. To test which time windows were significant, we also calculated, 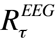, the correlation between Ŷ_τ_ and Y_τ_.

### Group level spatio-temporal analysis

For group level statistics, we first analyzed the 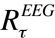 vectors across all subjects. The 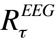 vectors were converted into their p-values, and for each time window (τ), used to compute combined Stouffer p-values ^66^. These group level results were then false discovery rate corrected (FDR) for multiple comparisons^67^. To identify group level voxels where our model predictions were significant, each subject’s p-value maps for the leave-one-out correlation were converted into their respective z-values, and voxel-wise significance was calculated using threshold-free cluster enhancement (TFCE) using a non-parametric randomization procedure implemented in FSL^68^. Voxels were considered significant if they passed a conservative false discovery rate threshold of p<0.01.

These significant voxels were then used as a mask to temporally localize activations by computing the voxels that were consistent in their direction ( positive (high stimulus evidence) or negative (low stimulus evidence)) and timing (τ window). To this end, we implemented a spatio-temporal TFCE (stTFCE) in both space (neighboring voxels) and time (neighboring time windows - response time window not included) and computed significance through a randomization procedure. 33000 permutations (1000 permutations per window) were run by randomly altering the sign of each subject and the temporal ordering of the windows, as we were testing whether the weights were consistent in sign, voxel space, and temporal window. P-values were calculated by comparing the true stTFCE value with the distribution of permuted values. Again, voxels were considered significant if they passed FDR correction at p<0.05 (high stimulus evidence: FDR-Corrected p<0.0019, low stimulus evidence: FDR-Corrected p<0.00036). Note, that now our number of multiple comparisons was the number of voxels in the FDR-mask (20256) times the number of time windows (33). We analyzed the response time separately with a standard TFCE randomization procedure implemented in FSL (Fig. S2D).

### Dynamic causal modeling

To validate the encoding model timing, we implemented single-state linear dynamic causal modeling (DCM) using DCM10 in SPM8 ^69^, and applied this to the BOLD data to test the hypothesis that the temporal sequence of BOLD activations we found in our EEG-fMRI encoding method was most likely, relative to other possible sequences of these same activations, given only the BOLD data. We used the results of the encoding model to select seven regions of interest that spanned the entire trial. For the first region (labeled 175 in our figures), we computed the union of activations during the 175ms and 200ms windows. Activations of the 225ms (225) and 250ms combined with 275ms (250) windows become the second and third regions. We computed the union of activations during the 325ms and 350ms windows to create the fourth (325). For the fifth region (400), we computed the union of the activations during the 400ms-450ms windows. For the sixth region (650), we computed the union of the activations during the 650ms and 675ms windows. Finally, the union of the activations during the 725-800ms windows was computed to create the seventh region (725). We removed any overlapping voxels between any of the regions and then extracted time series from individual subjects’ preprocessed functional data in MNI space by estimation of the first principal component within each region.

We constructed 11 models (Figure S6) to investigate the directed connectivity of these regions and validate the temporal ordering found by the encoding model. Each model was feed-forward with first node in each model as the input region. The first model was the temporal ordering of the regions inferred from our EEG-fMRI encoding model analysis. For five of the models, we randomized the temporal ordering of the early regions (175, 225, 250) and the late regions (325, 400, 650, 725) separately. For the other five models, we fully randomized the temporal ordering of all the regions.

We used fixed-effects Bayesian model selection (BMS) to compare these 11 models both on a single-subject level and at the group level. BMS balances model fit and complexity, thereby selecting the most generalizable model. It estimates the relative model evidence and provides a distribution of posterior probabilities for all of the models considered. We also compared families of similar models^70^; the model space was divided into two families (early/late or fully randomized).

### Drift Diffusion Model (DDM) and Confidence Proxy

The DDM models decision-making in two-choice tasks. Here, we treated the decision (correct vs. incorrect) as our two choices. A drift-process accumulates evidence over time until it crosses one of two boundaries (upper or lower) and initiates the corresponding response^68^. The speed with which the accumulation process approaches one of the two boundaries (a) is called drift-rate (v) and represents the relative evidence for or against a particular response. Recently, Philiastides, et al. ^45^ showed that for conditions in which the boundary (a) does not change, a proxy for decision confidence for each trial (i) can be computed by 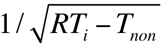.

We used Hierarchical Bayesian estimation of the Drift-Diffusion Model in Python (HDDM) to calculate the drift rate (v), decision boundary (a) and non-decision time T_non_ for each subject ^43^. Specifically, we modeled high and low stimulus evidence response time data separately. This was to ensure our confidence proxies were consistent within trial types. We included the response time and whether the subject got the trial correct. HDDM obtains a sequence of samples (i.e., a Markov chain Monte Carlo; MCMC) from the posterior of each parameter in the DDM. In our model, we generated 5000 samples from the posteriors, the first 1000 (burn-in) samples were discarded, and the remaining samples were thinned by 5%.

After modeling the DDM process, each trial’s (i) confidence proxy (CP) for each subject (j) was computed by 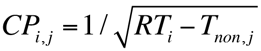 and then z-scored across trials where T_non,j_ was varied for high or low stimulus evidence trials, separately. Therefore, CP was a measure of relative trial confidence within difficulty levels.

### Confidence Proxy and Decision Replay

Trial to trial reactivation amplitude was defined as 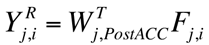 for each subject (j) and trial (i), where W_postAcc_ is the weight matrix from the encoding model thresholded by voxels that were significant in the group results from the 600-800ms windows. The mean of the 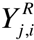 across time becomes a measure of “decision replay” strength for that trial (more negative y’s indicate more replay activation, more positive y’s indicate less replay activation). 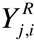 was quintiled for high and low stimulus evidence and the average confidence proxy was calculated within each quintile (Fig. 7). A linear mixed effects model^71^ was used to test if the slope of confidences across quintile grouping, 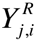, were significantly different from 0 while including stimulus evidence as a condition. Separate similar analyses with non-replay windows (175-250ms) and testing for behavioral accuracy were also performed (Fig. S7–8). To test the contribution of each cluster to the correlation with confidence, we implemented recursive feature elimination, where our features were clusters of significant voxels (> 48 voxels) during the 600- 800ms time window. This procedure removed clusters from the ‘replay’ network before calculating trial-to-trial reactivation. We then calculated the percent change in slope (reactivation x confidence proxy) when the cluster was removed compared to the total network. This procedure ranks cluster importance by sorting which clusters, when removed, had the strongest negative effect on slope height.

## Author Contributions

Conceptualization, J.M. and P.S.; Methodology, J.M., T.R.B., J.W. B.C., R.I.G. and P.S.; Investigation, J.M.; Software, J.M., B.C.;Writing - Original Draft, J.M. and P.S.; Writing - Review & Editing, J.M., T.R.B., R.I.G., J.W., and P.S. ; Funding Acquisition, P.S. ; Resources, J.M., T.R.B., J.W. B.C., R.I.G. and P.S.; Supervision, T.R.B and P.

## Acknowledgements

We would like to thank Jianing Shi for assistance in collecting the EEG/fMRI data. This
work was funded by National Institutes of Health Grant R01-MH085092, DARPA under Contract NBCHC090029 and the Army Research Laboratory and under Cooperative Agreement Number W911NF-10-2-0022.

